# The mechanical code of DNA links DNA bending nucleosome organization and codon choice

**DOI:** 10.1101/2025.09.09.675246

**Authors:** Aditi Biswas, Bailey Forbes, Jonghan Park, Aakash Basu-Biswas

**Affiliations:** Department of Biosciences, Durham University, Durham, DH1 3LE, United Kingdom; College of Medicine, Yonsei University, Seoul, Republic of Korea

## Abstract

Nucleosomes are the fundamental units of chromatin, yet how DNA sequence encodes the mechanics required for their formation remains incompletely understood. Using high-throughput measurements and predictive modeling, we show that nucleosomal DNA is distinguished not only by enhanced isotropic bendability but by coherent anisotropic bending aligned with the DNA helical repeat. Within nucleosome footprints, anisotropy vectors are phase-aligned across thousands of sequences, yielding ensemble-level signatures—such as ∼15 full rotations of bending direction—that directly mirror histone–DNA wrapping. We show that within nucleosomes, reinforcement of bending direction every helical repeat pre-disposes DNA to wrap smoothly around histones, lowering the energetic cost of nucleosome formation. Extending these principles to coding regions, we demonstrate that synonymous codon usage bias varies systematically around nucleosomes in a manner that optimizes DNA mechanical and structural features around nucleosomes and suggest that degeneracy of the genetic code has been exploited to encode both proteins and chromatin architecture. Together, our findings establish DNA anisotropy as a pervasive, sequence-encoded feature of nucleosomal DNA and identify codon usage as a key contributor to this hidden mechanical code.

## Introduction

Nucleosomes, formed by the wrapping of ∼147 bp of duplex DNA in ∼1.6 turns around an octamer of histones proteins, are the fundamental repeating subunit of chromatin in all eukaryotes. They play central roles in transcriptional regulation, genome accessibility, and higher-order compaction^1^.

It has long been recognized that mechanical directionality and energy cost in DNA bending is a fundamental determinant of how and where nucleosomes form and how DNA wraps around histones. Early structural studies have shown that nucleosomal DNA bends towards the minor grooves roughly every 10 bp along the nucleosome, stabilized by histone Arginine sidechains inserting into the minor groove^2,3^. This led to the hypothesis that sequence features within nucleosomal DNA may intrinsically favor smooth bending so that the minor groove faces inwards towards the histone core every 10.4 bp. Two classes of evidence supported this view. First, comparative sequence analyses revealed periodic enrichment of AA/TT/AT/TA (“WW”) dinucleotides at positions where structural models predict minor groove bending^4–7^. Second, biochemical and biophysical assays showed that “WW” motifs intrinsically facilitate bending towards the minor groove^8–10^.

Other studies extend these observations, implicating DNA mechanics, bending anisotropy, and shape as determinants of nucleosome organization. Local DNA shape such as minor groove width and intrinsic curvature strongly influence sequence-dependent nucleosome affinity, suggesting that sequence features might impact nucleosome formation via its impact on DNA bendability^11,12^. Anisotropic bending of DNA has been shown to facilitate nucleosome positioning and support the rotational positioning of DNA within nucleosomes^13,14^. Because many nucleosomes fall within protein-coding regions, it has also been speculated that **codon choice and nucleosome organization may have co-evolved**, with synonymous codon bias helping to accommodate mechanical constraints of chromatin packaging^15^.

Despite these insights, most evidence remains indirect. While high-throughput maps of nucleosome positioning exists, there exists significant gaps in our understanding of how sequence around nucleosomes impacts the biophysics of DNA. Periodicities in sequence motifs emerge only upon averaging across many nucleosomes and are largely absent in individual sequences ^16,17^. Likewise, models of nucleosome energetics rely heavily on structural inference or limited low-throughput assays ^18^. What has been lacking are direct, high-throughput measurements of DNA bending anisotropy and directionality within nucleosomes.

To address this gap, we recently developed loop-seq – a sequencing-based method to determine the local cyclization propensity of DNA in genome-scale throughput^17,19^. Initial analysis focused on the isotropic component of cyclizability, revealing that it peaks around known nucleosome dyads and dips in nucleosome depleted regions, thereby encoding a mechanical signature of nucleosome positioning. We also showed that these properties likely aid chromatin remodellers in properly positioning promoter-proximal nucleosomes.

Yet nucleosome bending in inherently anisotropic: the DNA must bend in specific directions, most notably into the minor groove every 10.4 bp. This raises two important questions: (i) is bending anisotropy indeed elevated in nucleosomal regions, and (ii) do bending directions indeed align in phase with the helical repeat? Loop-seq, by probing cyclization with three tethering geometries, implicitly encodes this information, though it was previously collapsed into the isotropic average. Building on this, we now leverage a machine-learning predictive framework to extract genome-wide anisotropy parameters directly from sequence. This allows us to test whether anisotropy is a pervasive, sequence-encoded feature of nucleosomal DNA and to explore how selection for isotropic and anisotropic DNA bendability has impacted the choice of codons.

## Methods

### Loop-seq, isotropic cyclizability, and anisotropy vector

Taking as input a chemically-synthesized library of up to ∼100,000 50 bp DNA fragments whose sequences are known and specified at will, loop-seq measured their intrinsic (isotropic) cyclizability in high-throughput^17,19,20^. However, to obtain isotropic cyclizability, cyclization propensity is measured for each fragment under three different tethering geometries. In the past^17^, we had assumed a simple sinusoidal model to extract a mean term from these three measurements of cyclizability, which we termed “Intrinsic cyclizability” (the isotropic component), or *C*_*0*_ (supplementary note 1). However, our model also produces anisotropy terms associated with cyclizability for every – a preferred direction of bending, *ϕ*, and an amplitude *A* (quantifying the extent to which bending is preferred in the ϕ direction (supplementary note 1)).

For every sequence, C_0_ represents its isotropic cyclizability, whereas (A, ϕ) together define an anisotropy vector, representing the preferred extend of DNA bending in a specific direction.

### Neural-nets based predictive model for c26, c29, and c31

We recently developed a neural-nets based predictive model to predict c26, c29, and c31 from any given input 50 bp DNA sequence, trained on the basis of loop-seq data^21^.

### Refined physical model for the dependence of C_0_, A, and ϕ on c26, c29, and c31

We discovered that our earlier model for the dependence of C0, A, and ϕ on c26, c29, and c31 introduced spurious noise into A at the frequency of the helical repeat, and its higher harmonics (Supplementary Note 2). We traced this noise to non-uniform scaling factors between these channels (Supplementary Note 2) and corrected for it by introducing a dependence of A and C_0_ on the tether location, in addition to tethering geometry specific dc offsets (Supplementary Note 3). The scaling parameters were trained on the basis of random DNA sequences, and applied uniformly to all future predictions.

## Results

### Isotropic and anisotropic DNA bending parameters around nucleosomes

We previously constructed a “Tiling Library”, comprising overlapping 50 bp fragments spanning from 600 bp upstream to 400 bp downstream of 576 randomly chosen yeast genes, with successive fragments offset by 7 bp^17^. Using loop-Seq we measured the cyclizability of these fragments in three specific tethering geometries, from which we extracted the intrinsic cyclizability C_0_ (the isotropic component of bendability), amplitude A, and phase ϕ (which together define the extent of anisotropic bending preference in a specific direction) of every fragment. Aligning sequences by the dyads of the +1 nucleosomes, the mean C_0_ profile (Fig. 1) shows the expected architecture: a marked dip over the upstream nucleosome-depleted region, a strong peak centered at the +1 nucleosome, and smaller peaks at +2 and +3 nucleosomes. This profile closely parallels independent nucleosome occupancy maps^22^, confirming that C0 reflects the general sequence preferences underlying nucleosome positioning.

**Figure 1.**
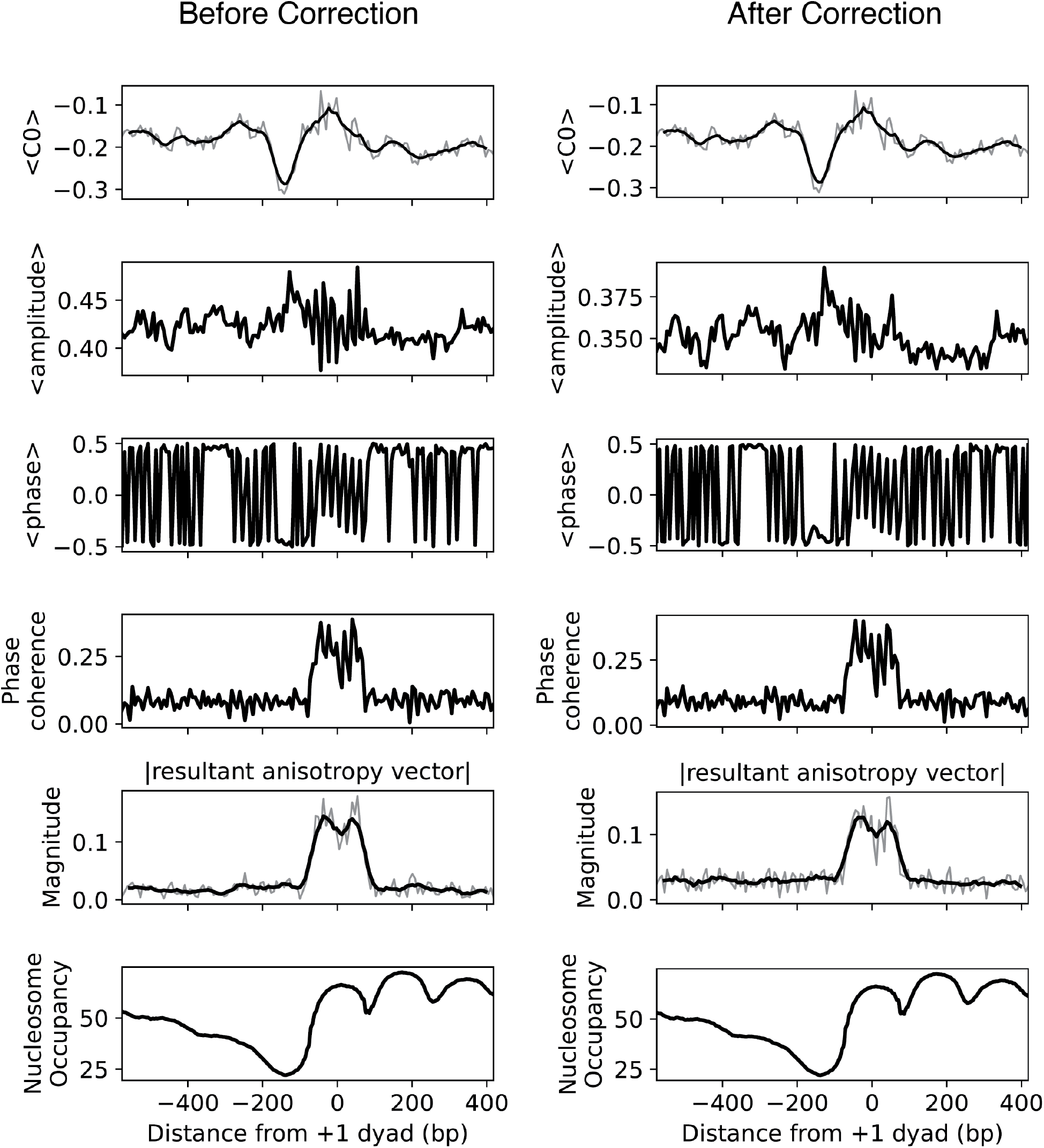
Mean C0, amplitude (A) and phase (f) as a function of position from the +1 nucleosome dyad, averaged over 573 DNA sequences spanning the +1 dyads of 573 randomly chosen genes in S. cerevisiae. Also plotted are the overall phase coherence of the angle of the anisotropy vector across different sequences, the magnitude of the vector average of individual anisotropy vectors across different sequences, and nucleosome occupancy, all as functions of distance from the +1 nucleosome dyad. See supplementary note 4 for plotting details. For each plot, quantities are plotted before and after scaling corrections (Supplementary notes 2-3).

Beyond isotropic flexibility, we examined anisotropy. The mean amplitude and circular mean phase exhibit pronounced oscillations within the 150-bp span of the +1 nucleosome but remain flat outside this region (Fig. 1). Far from the dyad, phase offsets across sequences are random and the circular mean of phase across all sequences is flat. Near the dyad, however, the oscillatory behavious in the circular mean of phase across hundreds of sequences imply phase coherence: bending directions add in register and rotate with the DNA helical repeat. Consequently, a direct measure of phase coherence across all DNA sequences as a distance *x* from the +1 nucleosome dyad, *R*(*x*) = |*e*^*iφ*(*x*)^|, is elevated only across the nucleosome footprint (Fig. 1). This shows that many different sequences share a common preferred bending direction at fixed distances from the dyad.

To capture direction and strength simultaneously, we computed the mean anisotropy vector

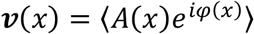

at every distance *x* from the +1 dyad, the mean being taken across all 573 sequences. We plot its magnitude |𝒱(*x*)| (Fig. 1) and find that across most positions, it is near zero, reflecting incoherent directions that cancel upon averaging. In sharp contrast, |𝒱(*x*)| rises prominently within the +1 nucleosome footprint, demonstrating that DNA in this region exhibit not only strong anisotropic bending preference, but also align their preferred bending direction coherently across many loci. These observations are consistent with a model where geometric constraints of nucleosome wrapping at successive superhelical locations have impacted the selection for phased bending anisotropy.

A caveat is that our 7-bp sampling introduces aliasing, precluding a precise estimate of the 10.4-bp periodicity from these data alone. Moreover, oscillations in the mean amplitude can partly arise from strong directional coherence in 𝜑 (coherent directions amplify any fluctuations at the helical frequency). These considerations motivated the analyses that follow, where we (i) increase positional resolution via the use of predictive models instead of directly measured data and (ii) reduced coherence-driven noise in amplitude.

To overcome the aliasing introduced by undersampling (which limits our ability quantify periodicities reliably), and to extend our analysis beyond the initial set of 573 genes, we decided to apply our previously developed neural network based model^21^ to predict c26 c29, c31 of any given 50 bp sequence, and subsequently to calculate C_0_, A, and ϕ. As a first step, we predicted the C_0_, A, and ϕ and used these values to reconstruct figure 1 (supplementary note 5) and recapture principal feature observed, suggesting that the neural nets model captured essential features of measured data.

However, when applying the model to totally randomly generated DNA sequences (as opposed to sequences aligned with respect to known nucleosomal dyads), we found that the reconstructed A (calculated from c26, c29, c31) contained strong spurious periodic components at the helical repeat (1×) and its first harmonic (2×) (Supplementary Note 2). We systematically traced these artifacts to imperfections in the forward–inverse mapping: scaling of the baseline term C_0_ or small d.c. channel offsets produce 1× ripples, while scaling mismatches in the amplitude term produce 2× ripples. To correct these biases, we trained on a large set of random DNA and optimized six global parameters (two C_0_ scalings, two A scalings, and two DC offsets) to minimize 1×/2× noise, followed by a single global 2× phase debiasing step (Supplementary Note 3). This calibration, once learned on random DNA, is then fixed and applied uniformly. We find that indeed the strong oscillations in A in Figure 1 are significantly suppressed, as expected for noise at the helical repeat adding in phase in the region of high phase coherence such as nucleosomes (Figure 1).

With this refined model, we then predicted C_0_, A, ϕ, phase coherence, and the magnitude of the mean anisotropy vector, across 9,702 annotated +1 nucleosomes of yeast^23^ (Pugh 2021). Each gene was tiled into overlapping 50 bp fragments offset by 1 bp, providing base pair resolution from -1,000 bp to +1,000 bp around the dyad. The resulting high-resolution, genome-wide profiles are shown in Figure 2.

**Figure 2.**
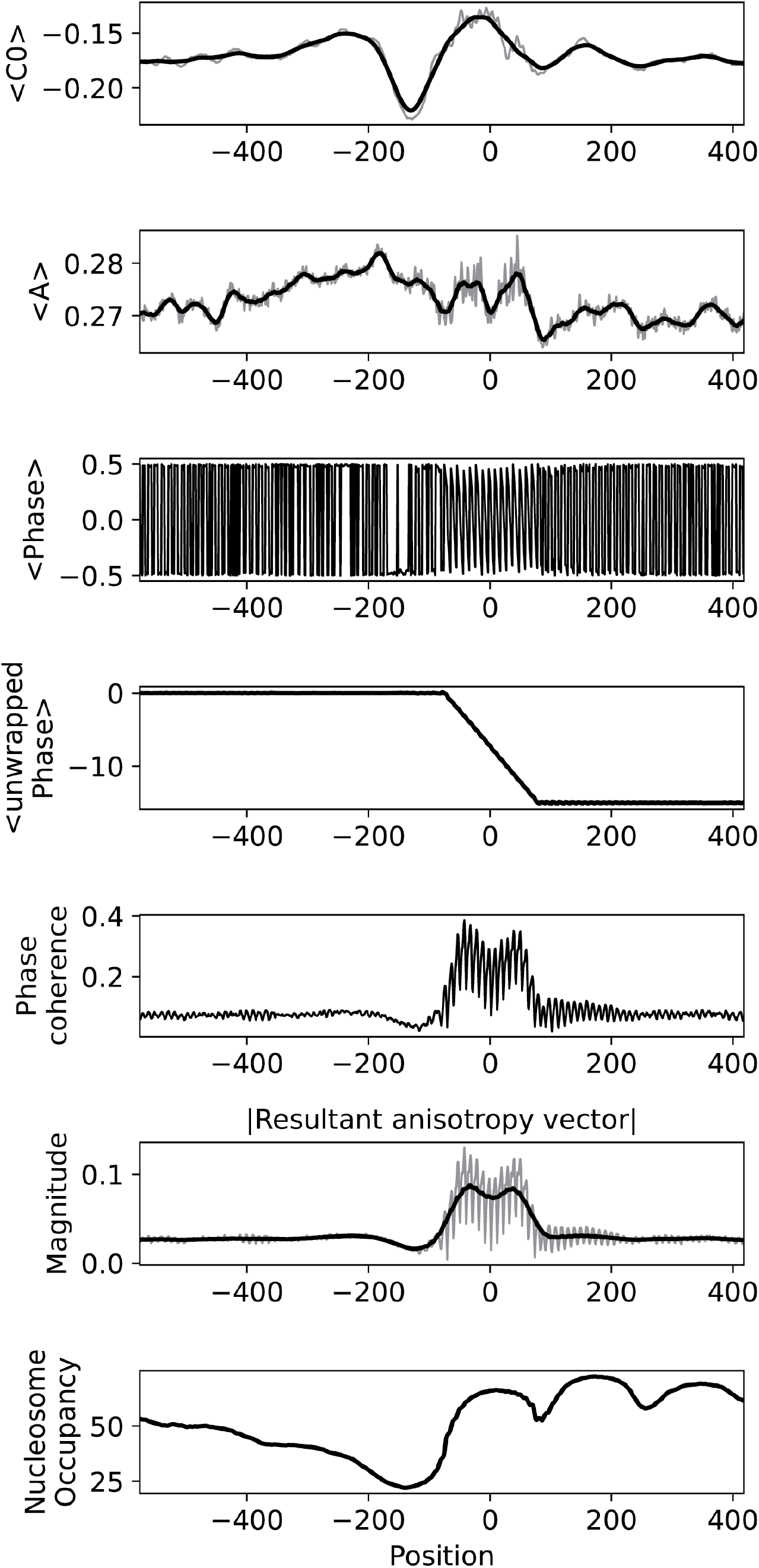
Mean C0, A, ϕ (wrapped and unwrapped), phase coherence, amplitude of vector mean of anisotropy vectors, and nucleosome occupancy, as functions of distance from the dyad of +1 nucleosomes.

The calibrated predictions substantially reduced noise in the amplitude channel. At the +1 nucleosome, we now observe two regions of elevated anisotropy flanking the dyad, suggesting that across thousands of promoters, DNA sequences have evolved to encode stronger bending anisotropy at these positions to facilitate the directional wrapping of DNA around the histone core.

The dense sampling also allows us to track the unwrapped mean phase angle 𝜑 as a continuous function of position. Strikingly, while the mean unwrapped phase angle is essentially flat across the genome but advances through ∼15 full rotations within the 150-bp footprint of the +1 nucleosome. This directly reflects the DNA helix wrapping around the histone core and mirrors the ∼14–15 helical repeats defined by successive minor-groove contacts. For any individual DNA sequence, the unwrapped phase inevitably advances at ∼10.4 bp per turn simply due to the geometry of B-form DNA (Supplementary Figure 2a in Supplementary Note 2). Consequently, ϕ along each of the 9,702 sequences spanning +1 nucleosome dyads follows its own trajectory, advancing 10.3 bp/turn starting from a different initial offset. In the absence of alignment these offsets cancel in the circular mean, producing a flat baseline outside the nucleosome footprint. Within the nucleosome, however, we find signature of strong alignment: φ trajectories are synchronized relative to the dyad, so that when averaged, the phase vectors add coherently, revealing the expected helical rotation even in the average vector. Thus, while φ advances in every sequence, only within nucleosomes are these rotations phase-aligned across molecules, yielding a clear collective signature of the nucleosome’s structural imprint.

To visualize this coherence directly, we examined the angular distributions of φ using polar histograms (“rose plots”) (Fig. 3). Outside the nucleosome footprint, these histograms are broad and stationary, showing no systematic change in orientation with position, consistent with phases being randomly offset across sequences. By contrast, within the 150-bp nucleosome span, the entire distribution rotates coherently with the DNA helical repeat, completing one full turn every ∼10 bp. This demonstrates that nucleosomes do not merely impose an average oscillation in the circular mean, but synchronize the bending directions of thousands of distinct sequences into a shared rotational register. The rose plots thus provide direct visual evidence that nucleosomal DNA exhibits population-wide phase alignment of anisotropy vectors, a property absent in non-nucleosomal regions.

**Fig 3.**
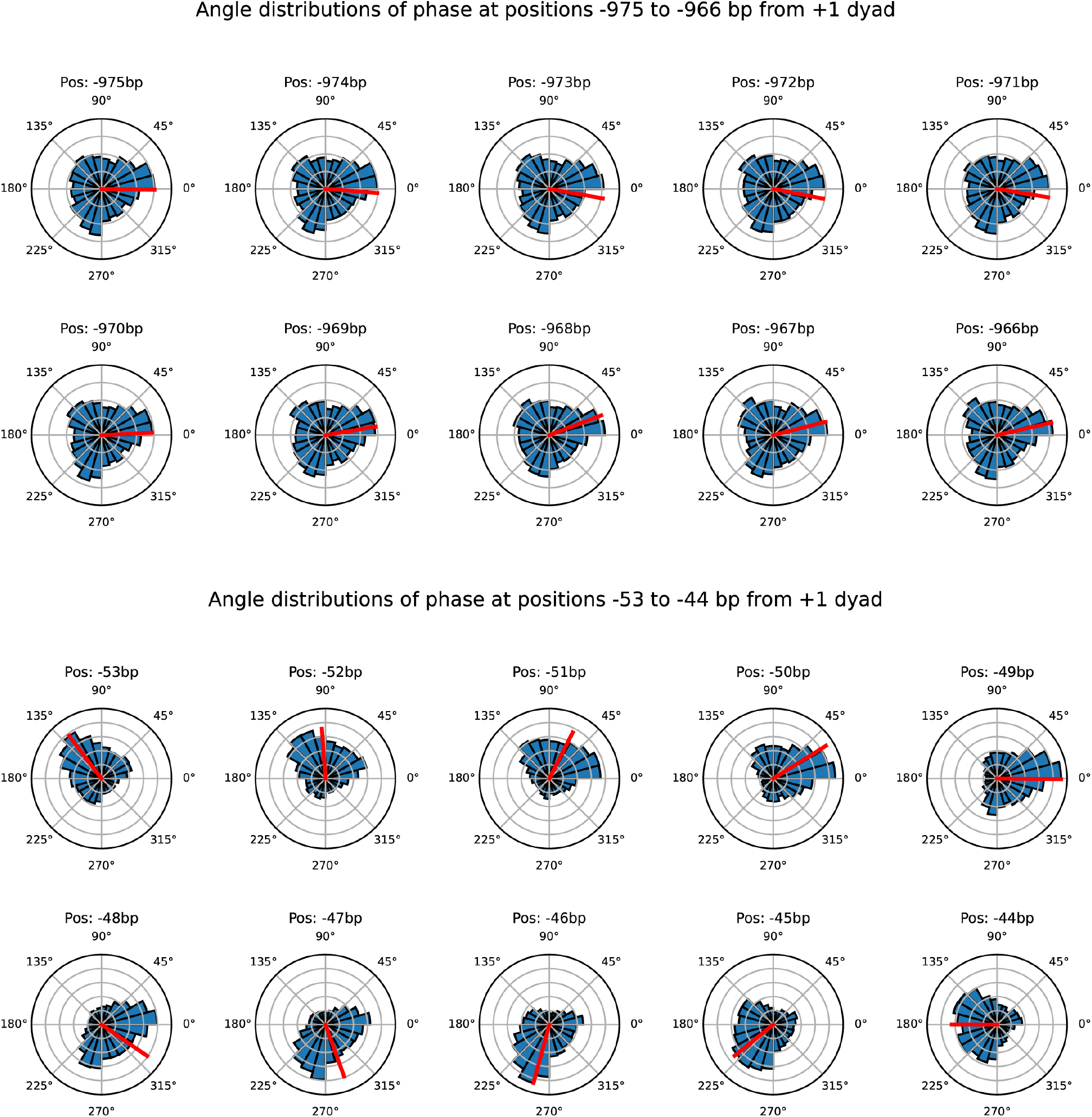
Polar histograms (rose plots) of ϕ, across 9,701 DNA sequences spanning +1 yeast nucleosomes. Histograms of phase angles across these sequences are plotted at a total of 20 locations (distances from the dyad): 10 locations (position -975bp to -966 bp) far from the dyad, and 10 locations (-53 to -44bp) close to the dyad.

The magnitude of the mean anisotropy vector of DNA (vector average of individual 9702 anisotropy vectors) at every position in the 2001 bp region around the dyad of the 9702 +1 nucleosomes is near zero everywhere other than in the 150 bp region around the dyad (Fig. 2). This mimics our direct measurements on a smaller subset of genes at lower resolution (Fig. 2d) and indicates phase coherence of the bending anisotropy direction in nucleosomal regions.

Moreover, within the footprint the mean anisotropy vector does not remain static: it rotates smoothly in a circle along the wrapped DNA (Fig. 4). Thus, the ensemble-averaged anisotropy vector directly tracks the periodic inward bending of the minor groove at successive superhelical locations.

**Fig 4.**
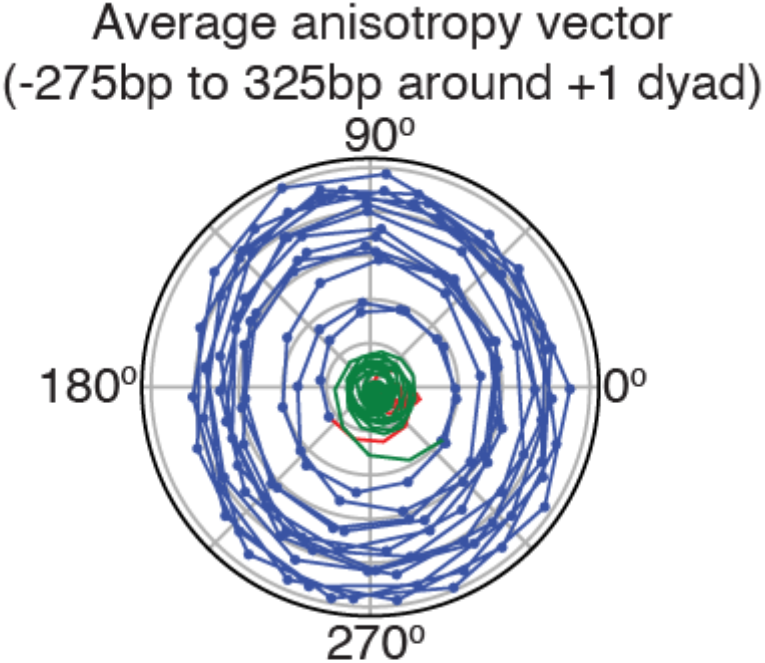
Tip of the mean anisotropy vector, averaged over 9,701 sequences, at various distances from the dyad. The vector is defined from the origin to the plotted point. Blue points refer to positions in the vicinity of the +1 nucleosome dyad (-75 bp to 75 bp), whereas red points lie further upstream (from -275 bp to -76 bp) and green points lie further downstream (76 bp to 325 bp). Points corresponding to neighbouring positions are joined by straight lines. Grid rings represent a magnitude difference of 0.02bp (a.u.).

Taken together, the analyses in Figs. 2–4 support a model in which sequence-encoded DNA bending anisotropy and its coherent alignment across molecules facilitate the periodic directional bending required for histone–DNA wrapping.

### Long-range phase correlation and bending coherence along nucleosomal DNA

If along a long DNA sequence the anisotropy amplitude is elevated and phase advances in register with the helical repeat, the DNA will intrinsically bend toward the minor groove once per turn, thereby favoring nucleosome wrapping. Because of the helical nature of DNA, phase values at adjacent positions are intrinsically correlated, and typically advances by ∼2π/10.4 bp. However, because our measurements of phase are defined over a 50 bp window, correlations are not expected to persist beyond ∼50 bp unless specifically encoded by the DNA sequence (as might happen within nucleosomes).

To test for such long-range phase correlations, we computed the average angular correlation of phase between positions separated by 100 bp. Across sequences aligned by the +1 nucleosome dyad, this correlation exhibits a sharp peak precisely at the dyad but is otherwise flat across the region (Fig. 5a, see Supplementary Note 6). This indicates that nucleosomal DNA sequences enforce directional bending anisotropy in register with the helical repeat, thereby stabilizing DNA wrapping at the dyad.

**Fig 5.**
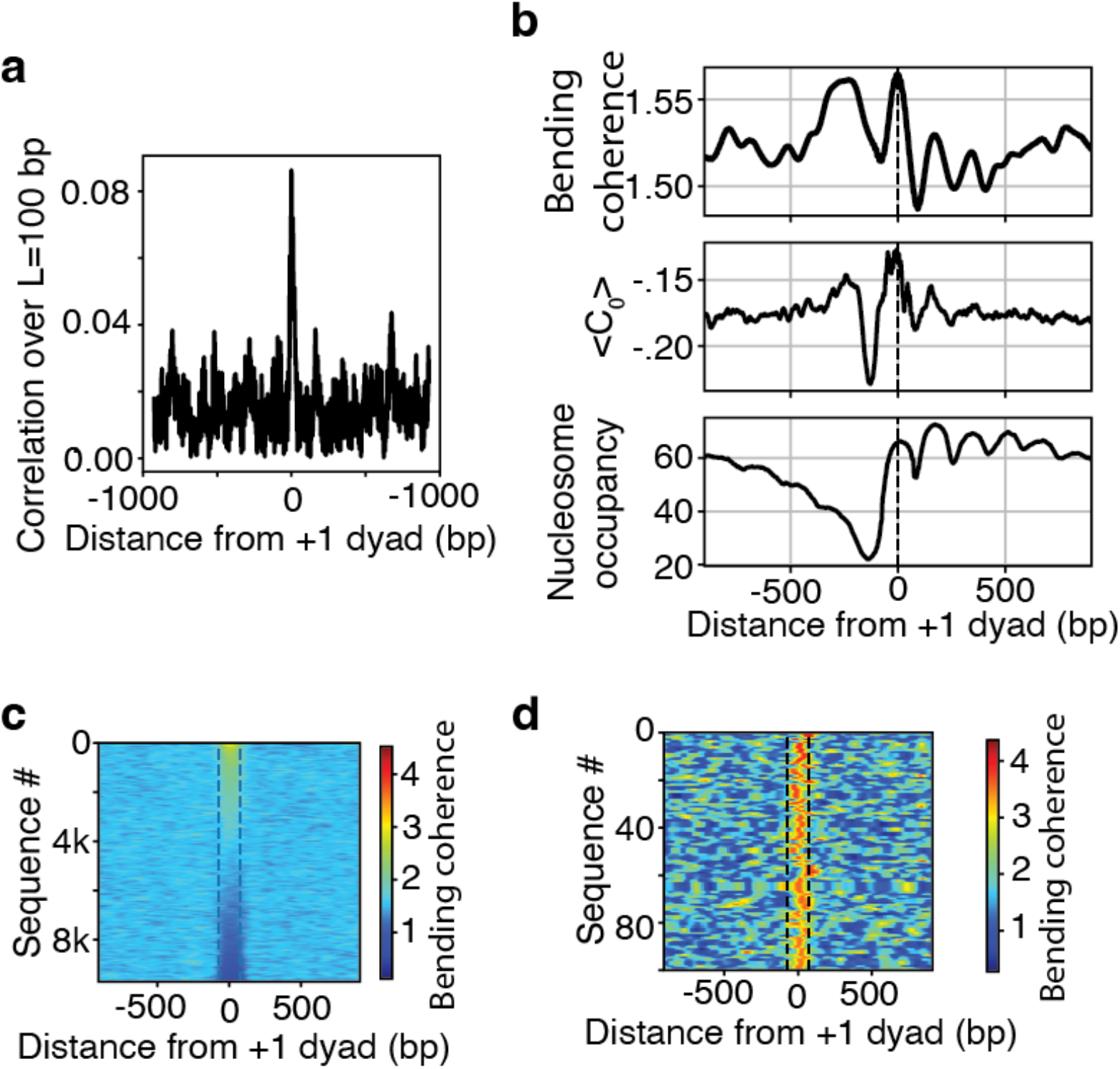
(a) Long-range phase correlation across nucleosomal sequences. For each of the 9,702 2001 bp DNA sequences spanning the +1 nucleosome dyads, the relative phase difference between positions j-50 and j+50 was calculated as 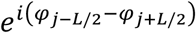, and these were averaged across sequences. The magnitude of this mean vector provides the phase correlation strength, and is plotted as a function of distance from the dyad. (b) Coherent bending signals revealed by phased vector summation. To test for helical-scale coherence, anisotropy vectors across all sequences were summed every ∼10.4 bp within a ±50 bp window centered on each position. The average magnitude of this summed vector is plotted as a function of distance from the +1 dyad. (c) Heatmap representing magnitude of the vector sum of the 10 vectors spaced ∼10.4 bp apart in a window around the position. Each row is a DNA sequence. Rows are ordered by the maximum signal in the vicinity of the +1 dyad. (d) Top 100 rows from c.

We have thus shown that nucleosomes are characterized not only by elevated anisotropy, but also by long-range correlation along nucleosomes, and coherence at any fixed distance from the dyad, across many nucleosomes. A key remaining question is whether bending anisotropy is reinforced every helical turn, as required for smooth nucleosome wrapping.

To address this, at any given distance from the +1 dyad, we summed the 10 anisotropy vectors surrounding that point, spaced roughly 10.4 bp from each other. If the sequence encodes a phasing of bendability in register with the DNA helix, then vectors separated by ∼10.4 bp will point in the same direction and add constructively. The magnitude of the summed vector, which we term ‘Bending Coherence’, therefore reports how well bending preferences reinforce across successive helical turns. When applied genome-wide, this analysis revealed striking peaks in bending coherence precisely at nucleosome dyads (Fig. 5b). In physical terms, the DNA sequence is pre-disposed to wrap smoothly around the histone octamer, lowering the energetic cost of nucleosome formation.

Visualizing bending coherence along individual nucleosomal sequences confirms that this effect is not driven by a few outliers: heatmaps reveal clear mechanical signature of high bending coherence around the vicinity of +1 dyads of a significant fraction of nucleosomes (Fig. 6c-d). Our observations provide a mechanistic explanation for how sequence features stabilize nucleosome positioning: DNA segments around dyads are encoded to bend in phase with the histone wrap.

**Fig 6a.**
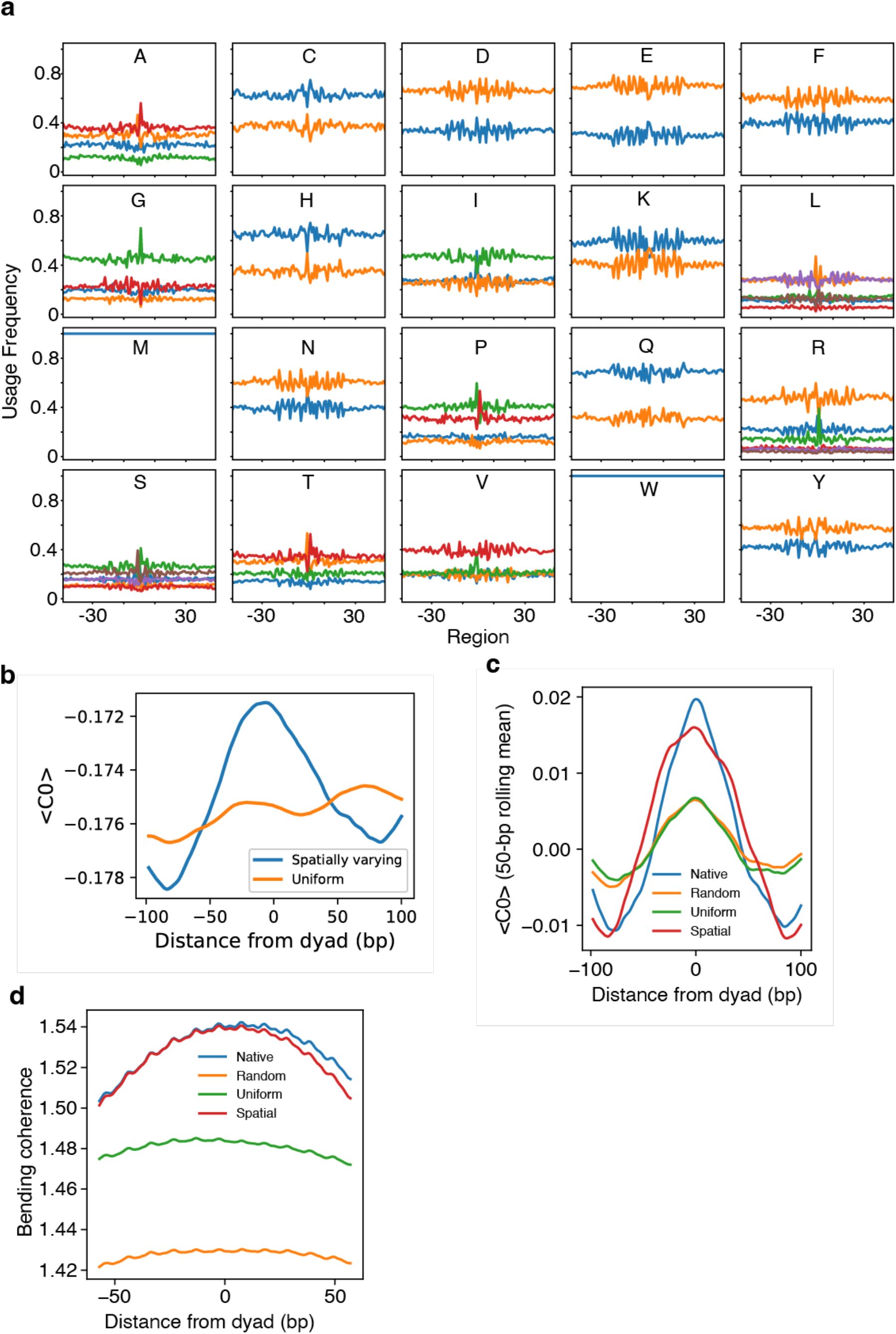
(a) Average usage frequency of individual codons as a function of distance from the dyads of gene-body nucleosomes. See supplementary note 7. Frequency is calculated from only amongst synonymous codons, and grouped according to encoded amino acid. (b) Mean intrinsic cyclizability (C0) as a function of position from the centre of 150,000 random 299 bp DNA sequenes, when these DNA sequences were generated by reverse translation from a common set of 150,000 99 amino acid peptides choosing codons that either reflect the uniform codon usage frequency in yeast (orange) or reflect how codon usage frequencies vary around the dyads of gene-body nucleosomes, assuming there is a nucleosome dyad at the centres of each fragmeng (blue). (c) <C0> as a function of distance from the dyad of gene body nucleosomes, for native DNA sequences (blue), and DNA sequences that have been condon randomized (but preserving the amino acid sequence) by choosing codons totally at random (orange), according to the genome-wise codon-usage frequency (green) and according to the observed variations in usage frequency of codons depending on distance from dyad, assuming a nucleosome dyad at the centre of the fragment (red).

### Codon Randomization

Our results thus far suggest that complex sequence patterns encode structural features within nucleosomal DNA that likely aid in the formation of nucleosomes. However, nucleosomal sequences within coding-DNA regions have to also encode for specific amino acid sequences. Degeneracy of the genetic code can in principle partially decouple amino acid sequence from the mechanics of the coding DNA. Therefore, we asked to what extent the choice of codons along genes has evolved to select for mechanical patterns that aid in nucleosome organization.

We first observed that codon usage frequencies are not constrant across the yeast genome: among synonymous codons, some are preferentially enriched near nucleosome dyads while others are depleted or shifted towards linker regions (Fig. 6a, Supplementary Note 7). In other words, codon usage varies systematically with distance from nucleosomes.

Why are certain synonymous codons found more frequently than others near nucleosomal dyads? To address this, we calculated the Bendability Quotient^19^ (BQ) of every codon (nucleotide triplet), which measured the average extent to which that codon, if embedded in any random 50 bp DNA sequence, would increase its isotropic flexibility (C_0_). We found that codons with higher BQ values are disproportionately enriched near nucleosome dyads: the codon usage frequency weighted mean BQ rises near nucleosome dyads, suggesting that synonymous codon choice contributes directly to flexibility in this region (Supplementary Note 8).

We next tested whether this spatial bias in codon usage frequency is sufficient, on its own, to generate nucleosome-like mechanical patterns. To do this, we generated random peptide sequences and reverse-translated them into DNA using codons sampled according to the observed position-specific codon usage frequencies around nucleosome dyads, assuming a nucleosome at the centre of the fragment. We found that this procedure reproduced a pronounced peak in mean C_0_ at the center of the sequences (Fig. 6b; Supplementary Note 8). Thus, the positional bias in codon usage is intrinsically capable of enhancing DNA bendability at nucleosome centers.

Finally, we asked whether native coding sequences actually exploit this mechanics. Examining gene-body nucleosomes, we found that real DNA sequences around dyads exhibit a local peak in both C_0_ and Bending Coherence (Fig. 6c–d; Supplementary Note 8). When the sequences were randomized by reassigning synonymous codons either completely at random, or uniformly, based on genome-wide codon frequencies, these peaks were diminished and the dyad–linker contrast was reduced. By contrast, when codons were resampled according to the observed spatially varying codon usage, the native profiles of C_0_ and Bending Coherence were closely recapitulated.

Together, these results demonstrate that codon usage bias is not only a product of translational optimization but also encodes structural features that facilitate nucleosome wrapping. Selection to maintain the mechanical requirements of nucleosomal DNA has therefore contributed to the evolution of codon choice, revealing an additional, structural layer of the genetic code.

## Discussion

Our results reveal a new layer of genome-encoded information: nucleosomal DNA is characterized not only by enhanced flexibility, but by coherent anisotropic bending aligned with the DNA helical repeat. Using loop-seq measurements and predictive modeling, we show, in high-throughput, that, for thousands of nucleosomes, DNA fragments centered on nucleosome dyads exhibit elevated anisotropy amplitude, synchronized phase trajectories, and long-range bending coherence. These findings establish DNA bending anisotropy as a pervasive, sequence-encoded property of nucleosomal DNA.

Nucleosomes enforce not just local anisotropy but collective phase alignment across molecules. While individual DNA sequences bend with the 10.4-bp helical repeat simply by virtue of B-form geometry, only within nucleosomes are these trajectories synchronized relative to the dyad. This coherence yields ensemble-level signatures—such as ∼15 full phase rotations across a nucleosome—that directly mirror the histone–DNA wrap. The rose plots provide striking visual confirmation: outside nucleosomes, anisotropy vectors are randomly oriented, but within the footprint, the entire distribution rotates coherently with the helix. This demonstrates that the structural constraints of nucleosome assembly are written into DNA sequence as population-wide mechanical order.

We also show that bending anisotropy is not confined to local motifs but reinforced over multiple turns of the helix. Peaks in long-range phase correlation and in “bending coherence” at nucleosome dyads reveal that DNA is encoded to wrap smoothly, lowering the energetic cost of histone association. This extends prior models of nucleosome energetics that emphasized anisotropic elasticity and twist–bend coupling^11,24,25^, now supported by genome-scale experimental data.

Our results also uncover a role for codon choice in encoding nucleosome mechanics. Codon usage frequencies vary systematically with distance from nucleosome dyads, and codons with higher bendability quotients are enriched near dyads. Randomization analyses confirm that these positional biases are sufficient to generate dyad-centered peaks in DNA bendability and coherence, even when amino acid sequences are held constant. Thus, degeneracy of the genetic code has been exploited to balance translational demands with the mechanical requirements of chromatin organization. This extends earlier speculation that nucleosome positioning signals could coexist with protein-coding constraints^15,26^ and provides direct mechanistic evidence for such dual encoding.

Together, these findings support a model in which DNA mechanics constitute a hidden but essential layer of the genome’s regulatory code. Nucleosome organization is guided not only by sequence motifs and remodeling factors but also by anisotropy vectors encoded in the DNA itself. Codon bias emerges as one mechanism by which coding regions reconcile protein sequence requirements with structural constraints on chromatin.

Looking forward, it will be important to ask how these signatures vary across species and chromatin contexts, and how they interact with nucleosome remodelers and transcription factors. In higher eukaryotes, where regulatory DNA is embedded in long coding and noncoding regions, similar mechanical encoding may shape nucleosome positioning on a larger genomic scale. More broadly, the ability to measure and predict DNA anisotropy at base-pair resolution offers a powerful tool for synthetic biology: engineering chromatin architecture by design of DNA mechanics.

In summary, this work establishes anisotropic bending as a genome-wide, sequence-encoded determinant of nucleosome organization and identifies codon usage as a key contributor to this mechanical code. By linking DNA mechanics, nucleosome architecture, and coding evolution, our results highlight the profound integration of the genetic and structural information within DNA.

## Author contributions

ABB designed research and supervised the project. AB and ABB performed all computation, analysis of SELEX data, and theoretical calculations. AB and ABB wrote the paper. BF performed nucleosome SELEX experiments. JP helped with cyclizability predictions. This work was supported by grant URF\R1\211659 to ABB and a Durham Doctoral Scholarship to AB. ABB is a Royal Society University Research Fellow.

## Notes

### Competing Interest Statement

The authors have declared no competing interest.

