## Supplementary Information for "The mechanical code of DNA links DNA bending nucleosome organization and codon choice"

### Supplementary Note 1: Loop-seq measures DNA bending in three distinct spatial directions

Loop-seq[1,2] quantifies the looping propensity of a 50 bp DNA fragment when it is tethered to a surface at three different points (26, 29, 31, referring to the distance in bp of the tethering point from the end of the DNA molecule). Thus, for any such fragment, loop-seq measured three numbers: c26, c29, c31, referred to the three cyclizability values.

Further, as our measurements require 50 bp fragments, we assign these values to the middle of the fragment. Therefore, if we tile a long DNA molecule, say of length 1000 bp, into 951 overlapping 50 bp fragments (fragment 1 from position 1-50, fragment 2 from 2 – 51, ..., fragment 951 from 951 – 1000), then c26, c29, c31 of the first fragment will be considered as the bending propensity of DNA in the three directions in space at position 25 along the parent sequence, and so on.

What do these three numbers c26, c29, c31 of a given 50 bp DNA fragment physically represent? We reason that each corresponds to the bending propensity of the DNA fragment in a different spatial direction (figure 1S). The biotin tether and bead surface provide steric hindrance to bending in one orientation, thereby favouring cyclization in the opposite direction. Figure S1 illustrates a cross-section through the midpoint of a 50-bp fragment, showing the position of the biotin tether relative to the bead surface.

This reasoning motivates our model:

$$C_n = C_0 + A\sin\left(\frac{2\pi}{10.4}n + \phi\right) \; for \; n = 26, 29, 31 \text{ ----------- (1)}$$

where $C_0$ is the intrinsic (isotropic) cyclizability of the DNA fragment, $A$ is the amplitude of anisotropy, reflecting the strength of preferential bending in a certain direction, and $\phi$ specifies the preferred bending direction in the DNA cross-sectional plane. If $A$ is zero, All $C_n$ are identical, as expected if the cyclizability of DNA is purely isotropic.

Therefore, we make three measurements of $C_n$ for n = 26, 29, 31, and solve the set of three equations (1) to obtain $C_0$, $A$, and $\phi$ for every 50 bp DNA sequence.

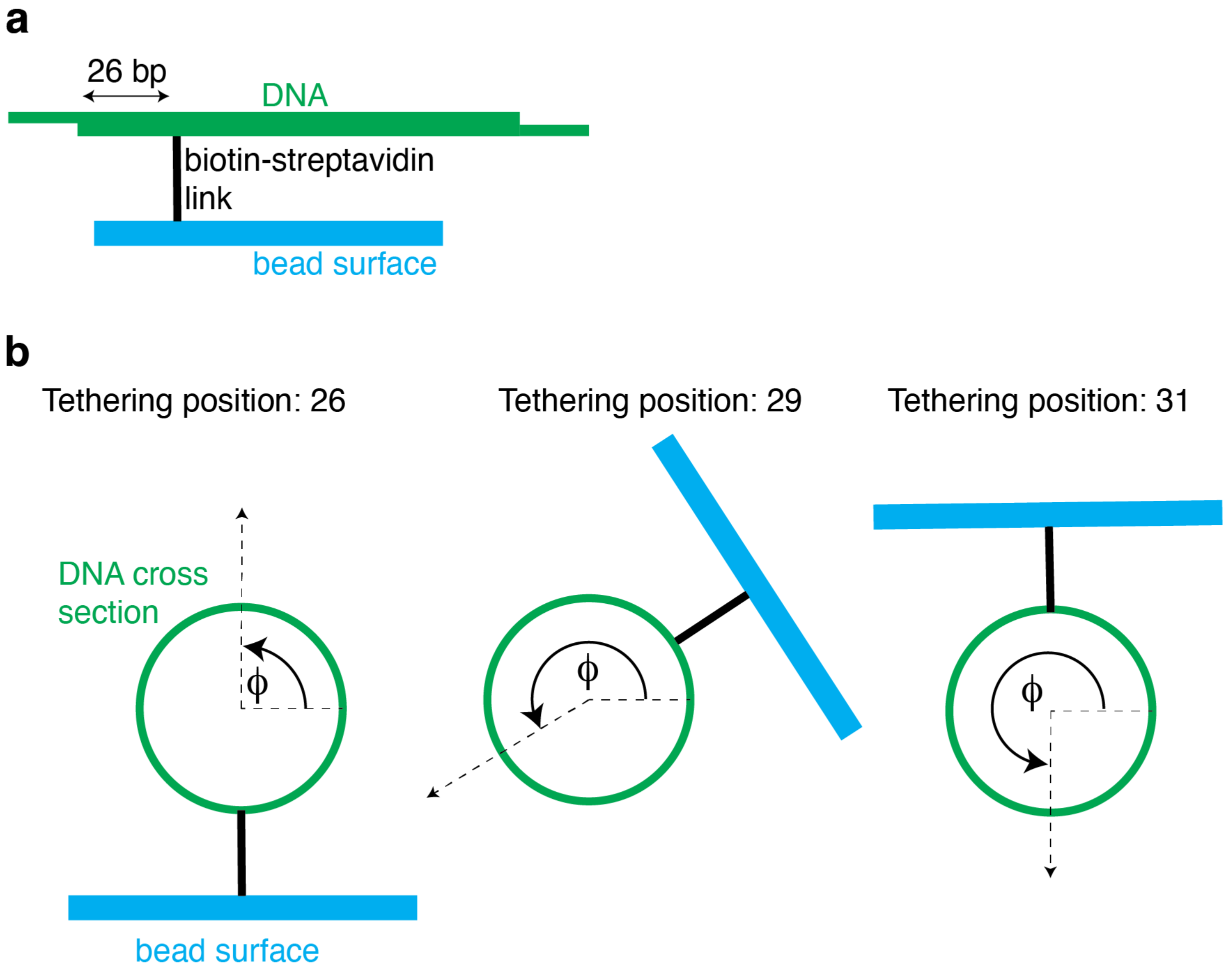


*Figure 1S: Cartoon representing DNA tethering in different geometries.* ***(a)*** *Basic geometry of the DNA cyclization assay. A DNA fragment tethered to a magnetic bead is permitted to cyclize via the annealing of complementary overahangs at either end. Loop-seq measures this cyclization propensity in high-throughput, for ~100,000 such DNA sequences at a time.* ***(b)*** *Cross sectional geometry. Green circle is the cross section of a 50 bp fragment emerging from the page. In cyan is the surface of the magnetic bead to which the fragment is tethered via biotin-streptavidin linkage (black solid line). We reason that the tethering geometry enforces a preferred direction of cyclization, namely "opposite" the biotin tether and bead surface. This direction can be defined by an angle $\phi$, with respect to some reference direction, and is indicated by the dashed arrow. The biotin is attached to some base that need not be at this cross-sectional plane shown. When the tethering point is moved 3 bp away (moving from the 26 to 29 tethering position), from the DNA's frame of reference, the bead/ biotin location has rotated by approximately* $\frac{2\pi}{10.4} \times 3 \sim 104^o$, *(and ~180°*

*when moved half the helical repeat away). Therefore, cyclizability measured by loop-seq at these three different tethering geometries (termed c26, c29, c31) reflect the propensity for the same 50 bp DNA fragment to bend in three distinct spatial directions.*

### Supplementary Note 2:

We first used a recently developed neural network model to predict c26, c29, and c31 from any given 50 bp DNA sequence[3] To test the model, we computationally generated 1,000 random DNA sequences of length 2,001 bp. Each sequence was parsed into 1,952 overlapping 50 bp fragments (fragment 1 = positions 1–50, fragment 2 = positions 2–51, etc.; see Supplementary Note 1). The neural nets model was applied to all fragments, yielding predicted values of c26, c29, and c31 for every position along every sequence. Each fragment was assigned to its midpoint (e.g. values for fragment 1 were placed at position 25 bp of the 2,001 bp sequence). Using the ideal model repreented by equation (1), we then solved for $C_0$, $A$, and $\phi$ for every fragment, assembling three matrices of size 1,000 × 1,952.

Because of the helical periodicity of DNA, we expect ϕ to advance by 2π / 10.4 ≈ 0.6 rad per base-pair step. As expected, the model reproduces this progression (Figure 2Sa).

Unexpectedly, however, the Fast Fourier Transform (FFT) of the plot of $A$ as a function of position along any sequence revealed strong periodic components at the helical repeat and its harmonics (Figure S1b). These periodicities are non-physical, as the input DNA sequences were completely random. We therefore systematically investigated the origin of this artifact in the $A$ term.

To do this, we artificially constructed 1,000 arrays of 2,001 elements each, each array hypothesized to reflect how $C_0$ varies with position along some hypothetical DNA sequence. Likewise, we generated arrays for A and phi for these 1,000 hypothetical DNA sequences. For each of 1,000 sequences, C0 was smoothly varied between −2 and 2 along the sequence, and A between 0.2 and 0.8. The phase was initialized at a random starting value per sequence and advanced by fixed increments of 2π / 10.4 rad per bp. From these synthetic inputs we generated c26, c29, and c31 using equation (1), and verified that our solver could correctly re-obtain $C_0$, $A$, and $\phi$ back.

We then introduced systematic distortions by applying scaling factors to test their effects. In effect, we tested what would happen if the tethering location impacted $C_0$, and $A$. First, we applied a scaling factor to the $C_0$ term for n = 29, 31 in equation (1). This multiplicative factor was 1.1, rendering our model:

$$C_{26} = C_0 + A sin\left(\frac{2\pi}{10.4} \times 26 + \phi\right)$$

$C_{29} = 1.1 \times C_0 + Asin\left(\frac{2\pi}{10.4} \times 29 + \phi\right)$

$C_{31} = 1.1 \times C_0 + Asin\left(\frac{2\pi}{10.4} \times 31 + \phi\right)$ -------------- (2)

In particular, we constructed $C_{26}$, $C_{29}$, $C_{31}$ from the initial $C_0$, A, ϕ using equation (2) above, but pretended we did not know about the existence of this scaling factor, and tried to use the ideal mode (equation (1)) to get back $C_0$, A, and ϕ. The reconstructed A exhibited spurious periodicity at the fundamental helical frequency (Figure S3a). Conversely, when a multiplicative scaling factor was applied to the *A* term (0.8 for $C_{29}$ and 0.7 for $C_{31}$) while constructing $C_{26}$, $C_{29}$, $C_{31}$, but ignored when attempting to solve back for $C_0$, A, and ϕ, the artifact shifted to the first harmonic (twice the helical frequency) (Figure S3b). In both cases, the true underlying signal contained no such periodicities, confirming that they arise from model mis-scaling rather than from the DNA sequences themselves.

Having demonstrated the causes of noise in the recovered amplitude A at the DNA helical frequency (1×) and its first harmonic (2×) via randomly generated data, we now mathematically formalize it below.

We consider three measured values $C_{26}, C_{29}, and\ C_{31}$ obtained from a 50 bp fragment and solve a linear 3×3 system per position to estimate (u, v), where $A = \sqrt{u^2 + v^2}$ and $\varphi = atan2(u, v)$. The fundamental period is P ≈ 10.4 bp, $k = \frac{2\pi}{P}$.

<u>1. Forward model and inversion</u>

Nominal forward model (per channel n ∈ {26,29,31}):

$C_n = C_0 + Asin(kn + \phi) = C_0 + u\ sin(kn) + v\ cos(kn)$
with $u = A.cos(\varphi)$ and $v = A.sin(\varphi)$. Stacking channels gives

$$\boldsymbol{C} = \boldsymbol{M}\ \boldsymbol{x}$$

Where $\boldsymbol{M} = \begin{bmatrix} 1 & sin(k.26) & cos(k.26) \\ 1 & sin(k.29) & cos(k.29) \\ 1 & sin(k.31) & cos(k.31) \end{bmatrix}$ and $\boldsymbol{x} = \begin{bmatrix} C_0 \\ u \\ v \end{bmatrix}$

We invert with the nominal M to obtain the measured $\boldsymbol{x}$ (called $\widehat{\boldsymbol{x}}$): $\widehat{\boldsymbol{x}} = \boldsymbol{M}^{-1}\boldsymbol{C}$

Once $\widehat{\boldsymbol{x}}$ is estimated, $\hat{A}$ and $\hat{\varphi}$ can be estimated as follows:

And, $\hat{A} = \sqrt{\hat{u}^2 + \tilde{v}^2}$, $\hat{\varphi} = atan2(\tilde{v}, \hat{u})$.

<u>2. Why $C_0$ scaling and offsets create 1x ripples in *A*</u>

Suppose the true C26, C29, C31 have a slightly different scaled contribution from C0: $C_n = \alpha_n C_0 + u.sin(k.n) + v.cos(k.n) + \gamma_n$. Then the true forward matrix is:

$$\boldsymbol{M'} = \boldsymbol{M} + \Delta\boldsymbol{M}, \qquad \Delta\boldsymbol{M} = \begin{bmatrix} \alpha_{26}-1 & 0 & 0 \\ \alpha_{29}-1 & 0 & 0 \\ \alpha_{31}-1 & 0 & 0 \end{bmatrix}$$

Therefore, the measured vector is:

$$\boldsymbol{C} = (\boldsymbol{M} + \Delta\boldsymbol{M})\boldsymbol{x} + \boldsymbol{d}$$

Inverting with the nominal **M**:

$$\widehat{\boldsymbol{x}} = \boldsymbol{M}^{-1}\boldsymbol{C} = \boldsymbol{x} + \boldsymbol{M}^{-1}\Delta\boldsymbol{M}\boldsymbol{x} + \boldsymbol{M}^{-1}\boldsymbol{d}$$

Now, $\Delta\boldsymbol{M}\boldsymbol{x} = C_0\Delta\boldsymbol{m_0}, \Delta\boldsymbol{m_0} = \begin{bmatrix} \alpha_{26}-1 \\ \alpha_{29}-1 \\ \alpha_{31}-1 \end{bmatrix}$

Hence,

$$\widehat{\boldsymbol{x}} = \boldsymbol{x} + C_0\boldsymbol{M}^{-1}\Delta\boldsymbol{m_0} + \boldsymbol{M}^{-1}\boldsymbol{d}$$

The term $C_0\boldsymbol{M}^{-1}\Delta\boldsymbol{m_0}$ is a constant vector in ($C_0$, u, v)-space proportional to $C_0$; its (u,v) components form a fixed bias vector **b**=($b_u$,$b_v$) that does not depend on $\varphi$. The DC offset adds yet another constant vector $\boldsymbol{M}^{-1}\boldsymbol{d}$ with the same property. On computing

$$\hat{A} = \sqrt{(u+b_u)^2 + (v+b_v)^2},$$

the cross terms $ub_u + vb_v = A\|\boldsymbol{b}\|cos(\varphi - \varphi_b)$ introduces a $cos(\varphi)$ modulation, i.e., a 1x ripple.

<u>2. Why *A* scaling mismatches create 2x ripples in *A*</u>

Suppose channels differ in their sin/cos gains (amplitude scaling): $C_n = C_0 + \beta_n u.sin(k.n) + \beta_n v.cos(k.n)$. Assuming $\beta_{26} = 1$ (and specifically, $\beta_{29} \neq 1$ and $\beta_{31} \neq 1$), then columns 2 and 3 of the matrix are perturbed:

$$\Delta\boldsymbol{M} = \begin{bmatrix} 0 & 0 & 0 \\ 0 & (\beta_{29}-1).sin(k.29) & (\beta_{29}-1).cos(k.29) \\ 0 & (\beta_{31}-1).sin(k.31) & (\beta_{31}-1).cos(k.31) \end{bmatrix}$$

Proceeding as above,

$$\widehat{\boldsymbol{x}} = \boldsymbol{x} + \boldsymbol{M}^{-1}\Delta\boldsymbol{M}\boldsymbol{x}$$

Because $\Delta\boldsymbol{M}$ multiplies only the (u,v) part, the (u,v) correction is linear in (u,v):

$\begin{bmatrix}\hat{u}\\ \hat{v}\end{bmatrix} \approx (\boldsymbol{I} + \boldsymbol{R}_2)\begin{bmatrix}u\\ v\end{bmatrix}$, $\boldsymbol{R}_2$ = lower right 2x2 block of $\boldsymbol{M}^{-1}\Delta\boldsymbol{M}$.

Writing $(u, v) = A(\cos\varphi, \sin\varphi)$,

$$\hat{A} = \|(\boldsymbol{I} + \boldsymbol{R}_2)A(\cos\varphi, \sin\varphi)\| = A\sqrt{a.\,cos^2\varphi + b.\,sin^2\varphi + 2c.\sin\varphi\cos\varphi}$$

Using the identities $cos^2\varphi = \left(\frac{1+\cos 2\varphi}{2}\right)$, $sin^2\varphi = \left(\frac{1-\cos 2\varphi}{2}\right)$, and $\sin\varphi\cos\varphi = \frac{1}{2}\sin 2\varphi$, this becomes:

$$\hat{A} \approx A(\kappa_0 + \kappa_c\cos 2\varphi + \kappa_s\sin 2\varphi)^{1/2}$$

giving rise to a 2x ripple.

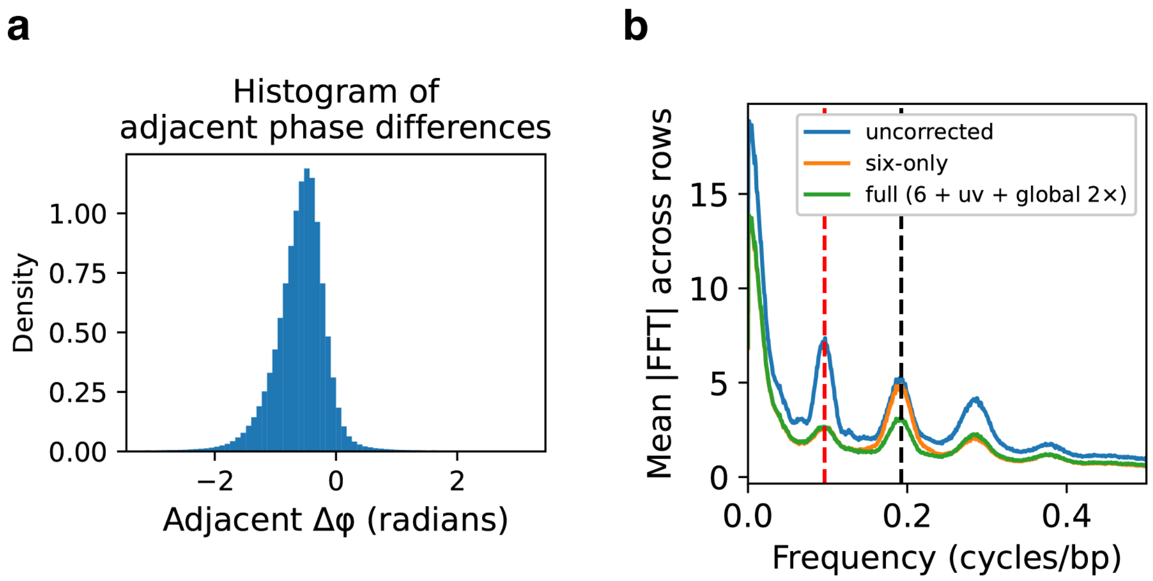


*Figure 2S: Analysis on random DNA sequences. 1,000 2,001 bp long DNA sequences were computationally generated randomly, parsed into 50 bp chunks, and passed through the predictive model to generate c26, c29, c31 as a function of position along the 1,000 sequences. This is described in Supplementary Note 2. (a) Histogram of $\Delta\phi$ between any two adjacent 50 bp fragments shows a strong peak at the expected value based on the helical repeat of DNA. (b) Noise in A. We first used the ideal model (1) to obtain C0, A, and f as a function of position along each of the 1,000 DNA sequences, from the predicted c26, c29, c31 values. We took a Fast Fourier Transform (FFT) of the A vs position plot of individual sequences, and averaged the 1,000 individual FFTs. The averaged power vs frequency plot is shown in blue, showing significant noise at the frequency of the helical repeat, and its harmonics. We then applied the six scaling factors trained on the basis on this dataset as described in supplementary note 4, which reduced some of these peaks*

*(orange curve). In addition, applying a 2x phase debiasing (supplementary note 4) reduced the noise even further.*

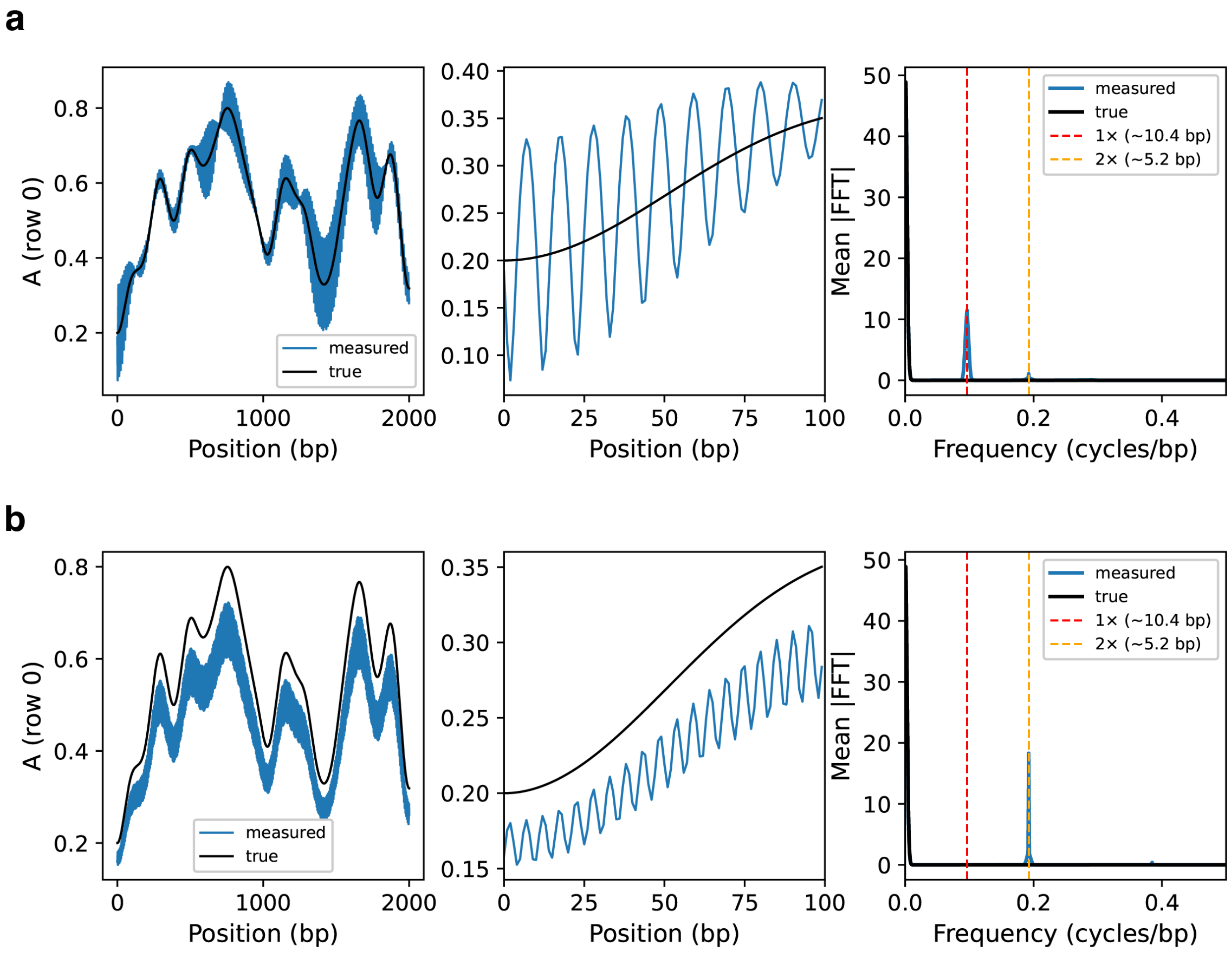


*Figure S3: (a) A vs position along one of the 1,000 DNA sequences, as hypothetically generated (black), and re-obtained by using the ideal model (equation (1)) in a situation where a scaling factor was introduced to scale the $C_0$ term in a tether location dependent manner. Shown also in a zoomen-in view of the first 100 basepairs, to make the noise at the helical period clearer. Plotted also in the mean of the individual FFTs of the 1,000 individual A vs position plots, showing the strong noise signal at the helical repeat (blue). The true signal (black) shows no such noise, as it was randomly generated. (b) Same as a, except multiplicative scaling was applied to the A term in a tether-location dependent manner.*

### Supplementary Note 3: Updates to the model and correction factors

The experiment generates $C_{26}, C_{29}, and\ C_{31}$. If we assume the ideal model (1) and try to generate $C_0$, A, and $\phi$, when the real physical model has scaling factors for A and $C_0$ terms, we demonstrated that we introduce spurious ripples in A (Supplementary Note 3). Therefore, we assume an updated model of the form

$$C_n = \alpha_n C_0 + \beta_n A sin\left(\frac{2\pi}{10.4} n + \phi\right) + \gamma_n \; for \; n = 26, 29, 31 \quad \text{-------------} \; (3)$$

$\alpha_n$: scale the $C_0$ column for channels 29, 31($\alpha_{26} = 1$)

$\beta_n$: scale sin/cos columns for channels 29, 31 ($\beta_{26} = 1$)

$\gamma_n$ : additive DC offsets to channels 29, 31 ($\gamma_{26} = 0$)

The above introduces six scalars. We learn these on random DNA sequences (no biology) by attempting to reduce the power peaks at 1x and 2x in the mean FFT signal in Figure 2Sb. Once learnt, these are simply 6 numbers that we freeze and use repeatedly when solving for $C_0$, A, and ϕ from any measured or predicted set of $C_{26}, C_{29}, and\ C_{31}$ values. To do so, we first mean center the c26, C29, C31 values, then apply the d.c. offsets $c_n{}' = c_n + \gamma_n \; for \; n = 29, 31$, and finally solve:

$$\begin{bmatrix} C_{26}{}' \\ C_{29}{}' \\ C_{31}{}' \end{bmatrix} = \begin{bmatrix} 1 & \sin(k.26) & \cos(k.26) \\ \alpha_{29} & \beta_{29} sin(k.29) & \beta_{29} cos(k.29) \\ \alpha_{31} & \beta_{31} sin(k.31) & \beta_{31} cos(k.31) \end{bmatrix} \begin{bmatrix} C_0 \\ u \\ v \end{bmatrix}$$

The parameters minimize mean row-wise FFT power of A of random DNA sequences at 1× and 2× with regularization.

<u>5. Residual 2x and global ϕ debiasing</u>

After the six-parameter fit, residual 2× often persists (amplitude still correlates with sin(2ϕ), cos(2ϕ)). We estimate a single global linear model on random DNA:

A ≈ b0 + b_s2·sin(2ϕ) + b_c2·cos(2ϕ)

and subtract the fitted 2× component everywhere:

$A_{corr}$ = A − (b_s2·sin(2ϕ) + b_c2·cos(2ϕ)).

We keep the intercept b0 intact (it sets the overall amplitude scale). We do not apply any global (u, v) offset here.

<u>6. Fully reproducible protocol (with guards)</u>

Inputs: random calibration set (c26_rand, c29_rand, c31_rand) and real set (c26_real, c29_real, c31_real). Shapes: R×C after reshape.

Preprocessing (both sets): row-center each channel (subtract row means).

Step 1 (learn 6 params on random): minimize $J = w1·P(1×) + w2·P(2×) + λ·‖θ−θ0‖^2 + λA·‖(sA−1)‖^2$, where P are band powers of mean row-wise $FFT(|A|^2)$.

Guards: bounds s0∈[0.6,1.6], sA∈[0.85,1.6], d∈[−1,1]; use Hann window in FFT; reject ill-conditioned solves (|det(M_6)| < 1e−12).

Step 2 (compute ϕ, A on random with learned 6 params).

Step 3 (global 2× model): fit [b0, b_s2, b_c2] by least squares of A on {1, sin(2ϕ), cos(2ϕ)} over all entries. Store β2_global = (b0, b_s2, b_c2).

Application to real data:

- Row-center channels; solve (u, v) with learned 6 params; compute A, ϕ.
- Apply global 2× debias: A ← A − (b_s2·sin(2ϕ) + b_c2·cos(2ϕ)) (keep b0).
- Use period P=10.4 unless otherwise stated; if columns are spaced by Δbp>1, convert FFT frequency to cycles/bp by dividing by Δbp.
- Sanity checks: report mean phase step per column ≈ 2π·Δbp/P; ensure A remains nonnegative; monitor 1×/2× band powers before/after.

7. Practical guards and limits

• Numerical stability: skip positions where |det(M_6)| < 1e−12; or fall back to pseudoinverse with Tikhonov damp (e.g., +1e−6·I).

• Regularization: keep sA near 1 to prevent amplifying 2×; penalize large offsets d29,d31; use bounded optimizers (e.g., L-BFGS-B).

• Windowing: use a Hann window before FFT; aggregate power in ±1 bin around 1×/2×; report both magnitudes and powers.

• Sampling: if columns represent Δbp > 1, use rfftfreq(..., d=Δbp) or divide frequencies by Δbp for cycles/bp.

• Period drift: if needed, re-estimate effective period by maximizing circular correlation of ϕ steps; otherwise keep P=10.4.

### Supplementary note 4: Plotting details for Fig. 1

The tiling library is identical to that described in supplementary note 9 of this[4] reference. Briefly 576 genes from *S. cerevisiae* were selected and the coordinates of the dyads of their +1 nucleosomes were noted. The 1044 bp DNA sequences from 601 bp upstream of the dyads (so position –601 if the dyad is defined as 0) to 442 bp downstream were obtained. Each of these sequences were tiled into overlapping 50 bp fragments- each fragment offset from the neighbor by 7 bp. For each gene, there were 143 such fragments:

Fragment 1: from –601 to –552
Fragment 2: from –594 to -545
...
Fragment 143: from +393 to +442

Therefore, across 576 such genes, there were 576 x 143=82,368 50bp fragments.

Loop-Seq was used measure the $C_{26}$, $C_{29}$, $C_{31}$ of these 82,368 DNA sequences, and either the uncorrected physical model (equation 1 in Supplementary Note 1) of the corrected model (equation (3) in supplementary note 3) were used to obtain $C_0$, A, and $\phi$ of these 82,368 sequences, and used to plot the first three panels.

To plot the magnitude of the vector average of the anisotropy vector, at each position, the individual 573 anisotropy vectors defined by A and f, were vector added and the resultant vector divided by 573. Plotted is the mean of this resultant vector as a function of distance from the dyad.

Phase coherence at each distance from the dyad defined as the magnitude of the resultant mean vector obtained by vector-summing unit vectors in the direction $\phi$ at every position from the dyad.

Nucleosome occupancy was obtained as described earlier[5,6].

### Supplementary Note 5: comparisons of measured vs predicted data

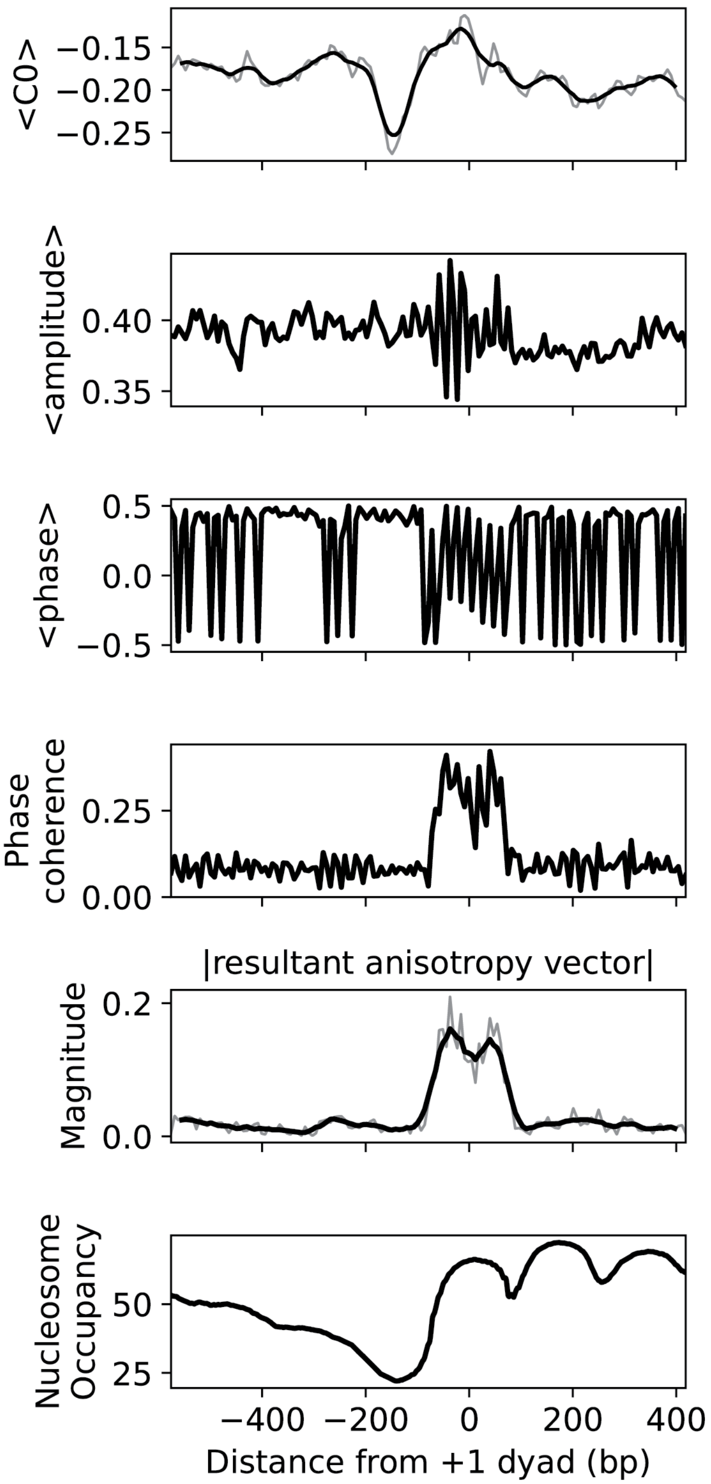


*Supplementary Figure 4S: Same as figure 1 (uncorrected), except c26, c29, c31 were obtained via predictions from the neural nets based model, as opposed to measured data.*

### Supplementary Note 6: Long-range phase correlation

To quantify long-range correlations in DNA bending phase, we analyzed sequences aligned at the dyads of +1 nucleosomes and computed the circular correlation of phase values across positions. Phase here refers to the angular direction of anisotropic DNA bending, defined modulo 2π, and is estimated locally over 50 bp fragments.

At an intuitive level, our goal was to measure whether the preferred bending direction at one site along the DNA is systematically related to the bending direction 100 bp downstream. Because phase is angular, standard linear correlation is inappropriate; instead, we use circular statistics based on complex unit phasors.

Formally, let $\varphi_{\{r,j\}}$ denote the measured phase (in radians, wrapped to [−π, π)) for sequence r at position j along the sequence. Here, we consider 9,702 sequences aligned to the dyad have obtained the phase at the 1,952 positions along each sequence, from position -975 to 975 bp with respect to the dyad of the nucleosome. We convert these to unit phasors:

$$z_{\{r,j\}} = e^{i\varphi_{\{r,j\}}}, \quad i^2 = -1$$

For a chosen lag L (here L=100 bp), we define the complex correlation at position j as:

$$c_j = \left(\frac{1}{N}\right)\sum_{i=1}^{N} z_{r,j-L/2} \cdot \overline{z_{r,j+L/2}}$$

where N is the number of sequences and the overbar denotes complex conjugation. This expression is equivalent to averaging the relative phase factor $e^{i\left(\varphi_{r,j-L/2}-\varphi_{r,j+L/2}\right)}$ across all sequences. The circular correlation magnitude as plotted in the magnitude of $c_j$, j ranging from -925 to 925 with respect to the nucleosome dyad.

### Supplementary Note 7: Identification of gene-body nucleosomes and plotting details for Fig. 6a

Identification of gene-body nucleosomes and the usage frequency of individual codons near their dyads:

The chromosome number, transcription start site (TSS), transcription termination site (TTS), and strand orientation of all 11,112 annotated yeast genes were obtained[6]. A comprehensive list of nucleosomal dyads was obtained from published maps[7]and genomic coordinates were converted to the SacCer3 assembly using liftover. From this list, we identified nucleosomes whose dyad-centered sequence of 297 bp (±148 bp around the dyad) lay completely within the coding region of an annotated gene (i.e. between the TSS and TTS). In total, 39,774 nucleosomes met this criterion. For each, the 297-bp sequence was extracted. When the associated gene was on the negative strand, the reverse complement of the sequence was used so that all sequences were oriented in the coding direction. Because codons are read in triplets, we required that the 297-bp sequence begin exactly at the first base of a codon rather than in the middle of one. To ensure this, we allowed the start of the 297-bp window to be shifted upstream by up to 2 bp if needed, determined from the known TSS of each gene, in order to maintain the correct reading frame. After this adjustment, each dyad-centered sequence corresponded to exactly 99 codons spanning the nucleosome. This list of 39,774 297 bp DNA sequences was noted and saved.

We now constructed a matrix called CODON_USAGE as follows: The matrix had 61 rows, each corresponding to the 61 non-stop codons, and 100 columns. Column 1 was names 'Codon', and was just the list of all 61 non-stop codons, grouped according to the amino acid they encode for. Columns 2 through 100 were named 'Region 1', …, 'Region 99', corresponding to the 99-codon region (297 bp region) straddling the dyads of gene-body nucleosomes. To populate this matrix, we followed the following process: For any given codon (AAA for example), the entry under the column 'Region 1' is simply the number of times the first three nucleotides in the list of 39,774 297 bp DNA sequences is AAA. It is, therefore, the counts of AAA in the first region, among the 99-codon region straddling the dyads of gene-body nucleosomes. Likewise, the entry under 'Region 2' is the number of times the second set of 3 nucleotides within the entries of the list matched 'AAA'. This procedure was followed to obtain the counts of every codon in all the 99 amino acid regions.

Fig. 6a: usage frequency of individual codons around the dyads of gene-body nucleosomes.

First we looked at the CODON_USAGE matrix and converted counts to usage frequencies as follows: At any region (i.e., for any column 'Region 1' through 'Region 99', we divided the counts by the sum of the counts of all its synonymous codons. This quantity was the usage frequency of that codon in that region, and by construction, the sum of the usage frequencies of all synonymous codons at each of the 99 regions is always 1. Plotted in Fig. 5a are the usage frequencies as a function of region number, of all 61 non-stop codons. Region number has been converted to DNA coordinates (as opposed to regions 1-99) by first multiplying by 3 and then offsetting to make the centre of the region (i.e. the nucleosome dyad) equal to 0. Synonymous codons are grouped together in individual sub-panels, and the amino acid they encode for has been noted. Therefore, for any amino acid and at any position, the usage frequencies of one of one of its possible synonymous codons reflect the probability that were the amino acid to be encoded by DNA at that distance from a nucleosome dyad, it would be encoded by that specific synonymous codon rather than by other synonymous codons.

### Supplementary Note 8: why are certain codons more prevalent at nucleosome dyads?

Overview:

For every amino acid, why has the yeast genome evolved to use certain synonymous codons more frequently than others near nucleosomal dyads (as seen in Fig. 6a and explained in

Supplementary Note 7)? Do these codons, on average, make surrounding DNA more flexible? To answer this question, we first measure the Bendability Quotient[4] (BQ, see below) of every trinucleotide (i.e., codon), which reflects the average propensity of that trinucleotide to contribute to the cyclizability of any random 50 bp DNA sequence within which it may be present. Then, for every amino acid, we calculated the average codon usage frequency of all its synonymous codons at each of the 99 positions straddling the dyads of the gene-body nucleosomes. Finally, we plotted, for every amino acid, the codon-frequency weighted mean BQ of all the possible synonymous codons at every position in this 99-codon region (see below). We indeed show that this mean BQ rises near the dyad in the case of several amino acids, suggesting that near nucleosome dyads, for every amino acid, those codons which have higher BQs occur more frequently (Supplementary Note 7).

We next generated a totally random set of 150,000 99 amino acid long peptide sequences and reverse translated them back to a set of 150,000 297 bp long DNA sequences. For each peptide, codons were chosen among possible synonymous codons, with probabilities given by the observed codon usage frequencies of these synonymous codons in that region (Fig. 6a, also described in supplementary note 7). We then parsed each DNA sequence into a set of 248 overlapping 50 bp fragments, each offset from the previous by 1 bp. We predicted C0 of each of these fragments using our neural nets based model. The C0 value of the first such 50 bp fragment of a 297 bp sequence was assigned to represent C0 at a position 25 bp from the left edge of that DNA fragment (or position -123 bp with the centre of the DNA being the 0 mark). Plotted in Supplementary Figure 5b is the mean C0 as a function of position from the centre of the fragments, averaged over all fragments. We see a peak in C0 near the middle of the fragments, suggesting that even for totally random amino acids, this particular form of spatial variations in codon usage bias is enough to bias the centre of the coding DNA fragment to have higher C0.

We then took the same set of 150,000 random 99-amino-acid protein sequences and reverse-translated them into DNA, but this time codons were chosen independently at each position according to the overall codon frequencies, without regard to position. The overall codon usage frequency of a certain codon is essentially the ratio of the sum of its counts in the entire matrix CODON_USAGE (across all 'Region' columns) divided by the sum of the counts of it and all other codons that are synonymous to it). Predicting C0 in a similar fashion yielded a flat profile without a dyad-centered peak.

Calculating bendability quotients:

Bendability quotients of all 64 trinucletides (i.e. codons) were calculated from measured intrinsic cyclizability values of 12,472 randomly chosen genes in *S. cerevisiae*. This mimics what we had done earlier in the case of di- and tetra-nucleotides[4]. Shown below are the bendability quotients of the 64 codons:

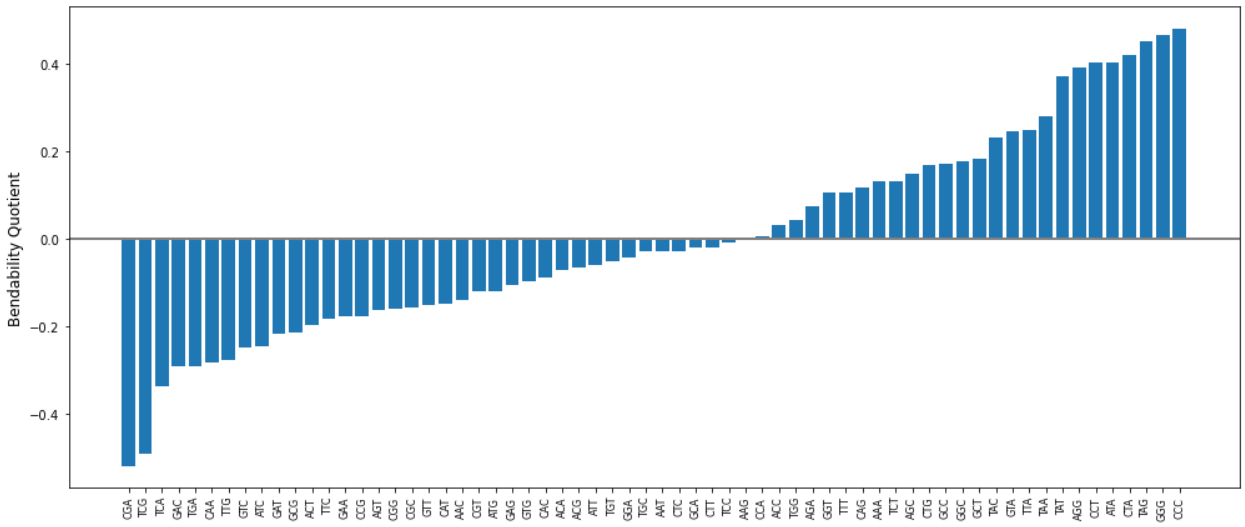


<u>Plotting of codon frequency-weighted mean BQ of all non-stop codons:</u>

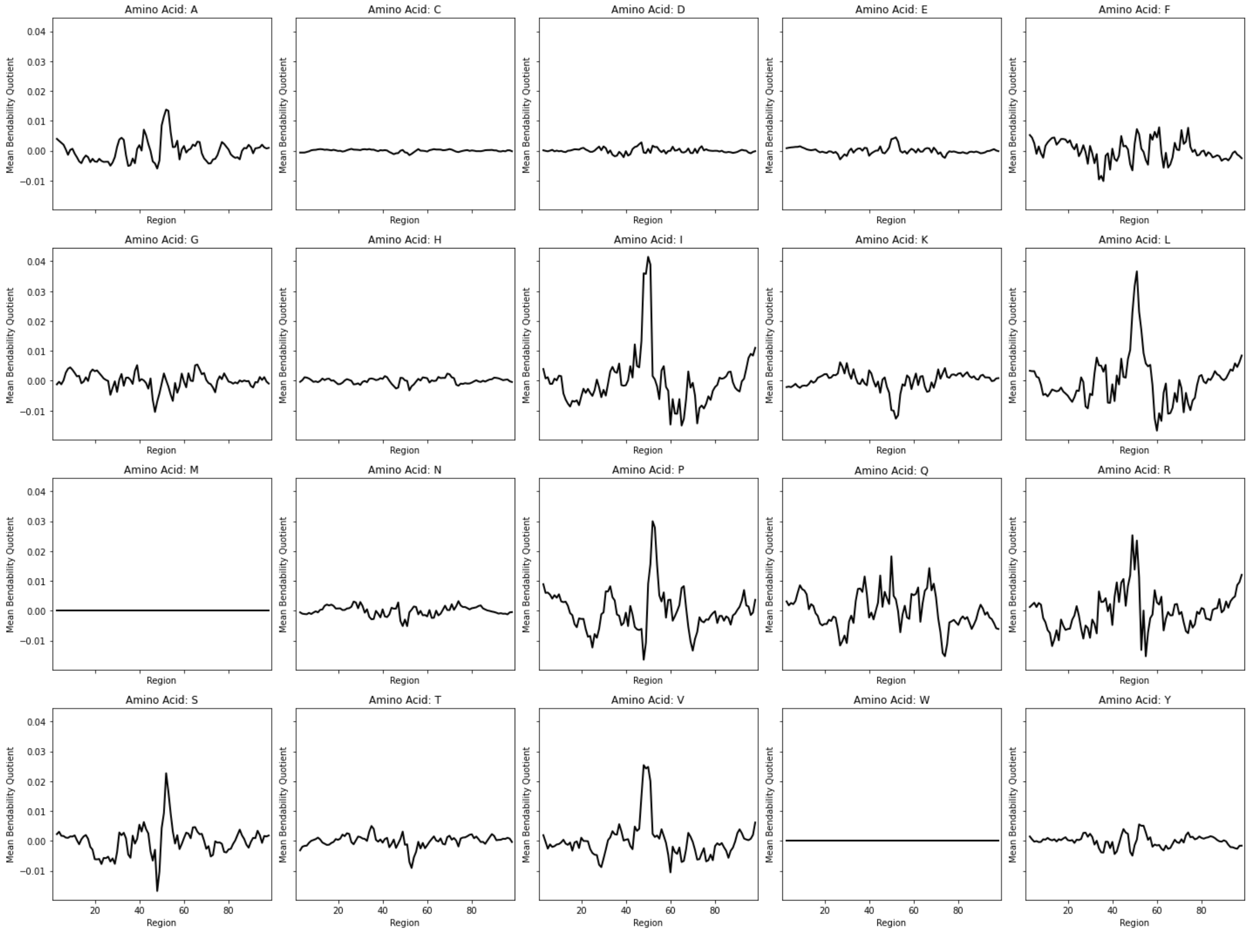


*Figure 6S: Codon usage frequency weighted mean bendability quotient among synonymous codons, around gene-body nucleosome dyads.*

### **<u>Supplementary Note 9</u>**

We investigated how this variation in codon usage frequency around dyads of coding nucleosomes in yeast has shaped the mechanical profile of nucleosomal DNA. From the 39,774 gene-body nucleosomes we identified in yeast, we created 6 lists (L1 – L6) of 39,774 297 bp DNA sequences. L1: the native 39,774 297 bp DNA sequences around these dyads, L2: a codon-randomized version of L1 that preserves the amino-acid sequences encoded, but codons are chosen totally at random from among possible synonymous codons with equal probability, L3: a uniform bias codon randomised version, where for every amino acid, a codon is chosen from among the possible synonymous codons with a probability that reflects the overall codon-usage frequency of all codons (as done in fig. 6b), and L4: a spatially-varying codon randomized version where the probability of choosing synonymous codons depends on the position of the amino acid along the 99 aa fragment (Fig. 6a). We further created two additional lists of 39,774 297 bp DNA sequences – L5: a totally random set of DNA sequences that only preserves the same average profile of GC content along the 297 bp region as list L1, and L6: a totally random list of 39,774 297 bp DNA sequences. L5 and L6 do not encode for the same peptide sequences, while L1-L4 all do (and match the native peptide sequence in yeast).
